# Saccade-responsive visual cortical neurons do not exhibit distinct visual response properties

**DOI:** 10.1101/2022.11.21.517415

**Authors:** Chase W. King, Peter Ledochowitsch, Michael A. Buice, Saskia E. J. de Vries

## Abstract

Rapid saccadic eye movements are used by animals to sample different parts of the visual scene. Previous work has investigated neural correlates of these saccades in visual cortical areas such as V1, however how saccade-responsive neurons are distributed across visual areas, cell types, and cortical layers has remained unknown. Through analyzing 818 one-hour experimental sessions from the Allen Brain Observatory, we present a large-scale analysis of saccadic behaviors in head-fixed mice and their neural correlates. We find that saccade-responsive neurons are present across visual cortex, but their distribution varies considerably by transgenically-defined cell type, cortical area, and cortical layer. We also find that saccade-responsive neurons do not exhibit distinct visual response properties from the broader neural population, suggesting the saccadic responses of these neurons are likely not predominantly visually-driven. These results provide insight into the roles played by different cell types within a broader, distributed network of sensory and motor interactions.

**Highlights:**

- Saccadic eye movement behaviors in head-fixed mice tend to occur in bursts, preferentially along the horizontal axis, and do not strongly depend on visual stimulus.
- Distributions of saccade-responsive neurons vary considerably by transgenically-defined cell type, visual area, and cortical layer. They are most prevalent in dorsal visual areas AL/PM/AM, inhibitory neurons, and deeper cortical layers.
- The majority of saccade-responsive neurons are selective for saccades in a particular direction, with an overwhelming preference for temporal over nasal saccades.
- Saccade-responsive neurons do not exhibit distinct visual response properties, suggesting saccade neural responses are not likely to be predominantly visually-driven.

## Introduction

Saccades are rapid eye movements that quickly shift the visual field between two points of fixation. Unlike primates, mice have no fovea, yet numerous studies have previously found that mice make saccades in both freely moving and head-fixed contexts (Meyer et al., 2020; Sakatani & Isa, 2007; Samonds et al., 2018). In the head-fixed context, these saccades are almost exclusively conjugate—both eyes moving in unison in the same direction—and occur preferentially along the horizontal direction (Meyer et al., 2020; Samonds et al., 2018). It has also been previously established that neurons in primary visual cortex (V1) modulate their responses to these saccadic eye movements (Itokazu et al., 2018).

Recent experimental findings in mice implicate the lateral posterior (LP) nucleus, a higher-order thalamic nucleus that is the rodent homolog of the thalamic pulvinar nucleus, as the primary non-visual source of saccade signals in cortex (Roth et al., 2016; Miura & Scanziani, 2022). In V1, LP projects to layers 1 and 5, and in higher visual areas the projections are to layers 1 and 4 (Herkenham, 1980). Recent electrophysiological work has found that this LP saccade signal encodes saccade direction and contributes to a subset of V1 neurons responding to eye movements in a direction-selective manner (Miura & Scanziani, 2022).

While it has been well-established that saccade-responsive neurons exist in visual cortex, it remains unknown, however, how these saccade-responsive neurons are organized. In the present study, we seek to address: (1) how saccade-responsive neurons are distributed across different visual areas, cortical layers, and genetically-defined cell types; and (2) how visual response properties of these saccade-responsive neurons compare to the visual response properties of the broader neural population. Specifically, we examined the saccadic eye movement behaviors across transgenic lines of mice and characterized the saccade response properties of different types of cells across visual areas and cortical layers. We analyzed physiological data from the Allen Brain Observatory Visual Coding dataset (publicly available through brain-map.org and the AllenSDK Python package), which consists of calcium imaging recordings in head-fixed mice across 6 visual areas, 4 cortical layers, and 14 transgenically-defined cell types in the mouse visual cortex (Table 1; *Allen Brain Observatory -- 2-Photon Visual Coding*, 2016; de Vries et al., 2020). We found saccade-responsive neurons distributed across all visual cortical areas, layers, and cell types within this dataset. However, this distribution was not uniform, as we observed comparatively more saccade-responsive neurons in certain classes of inhibitory interneurons and in deeper cortical layers. Furthermore, we found that these saccade-responsive neurons do not have distinct visual response properties, suggesting that saccade responses are not a uniquely visually-driven response.

**Table 1:** Transgenic line abbreviations

| Abbreviation | Full Transgenic Cre Line | Number of Mice | Number of Sessions | Number of Neurons Total | Number of Neurons Analyzed |
| --- | --- | --- | --- | --- | --- |
| **Emx1;Ai93** | Emx1-IRES-Cre; Camk2a-tTA; Ai93(TITL-GCaMP6f) | 17 | 83 | 8866 | 5020 |
| **Slc17a7;Ai93** | Slc17a7-IRES2-Cre; Camk2a-tTA; Ai93(TITL-GCaMP6f) | 24 | 124 | 8512 | 7707 |
| **Slc17a7;Ai94** | Slc17a7-IRES2-Cre; Camk2a-tTA; Ai94(TITL-GCaMP6s) | 1 | 6 | 817 | 817 |
| **Cux2;Ai93** | Cux2-CreERT2; Camk2a-tTA; Ai93(TITL-GCaMP6f) | 32 | 121 | 10181 | 6832 |
| **Rorb;Ai93** | Rorb-IRES2-Cre; Camk2a-tTA; Ai93(TITL-GCaMP6f) | 18 | 61 | 4738 | 3485 |
| **Scnn1a;Ai93** | Scnn1a-Tg3-Cre; Camk2a-tTA; Ai93(TITL-GCaMP6f) | 6 | 21 | 1513 | 935 |
| **Nr5a1;Ai93** | Nr5a1-Cre; Camk2a-tTA; Ai93(TITL-GCaMP6f) | 20 | 71 | 2122 | 1353 |
| **Rbp4;Ai93** | Rbp4-Cre_KL100; Camk2a-tTA; Ai93(TITL-GCaMP6f) | 21 | 72 | 1649 | 1399 |
| **Fezf2;Ai148** | Fezf2-CreER; Ai148(TIT2L-GC6f-ICL-tTA2) | 8 | 25 | 1346 | 1338 |
| **Tlx3;Ai148** | Tlx3-Cre_PL56; Ai148(TIT2L-GC6f-ICL-tTA2) | 3 | 12 | 979 | 979 |
| **Ntsr1;Ai148** | Ntsr1-Cre_GN220; Ai148(TIT2L-GC6f-ICL-tTA2) | 9 | 33 | 1180 | 1180 |
| **Sst;Ai148** | Sst-IRES-Cre; Ai148(TIT2L-GC6f-ICL-tTA2) | 25 | 91 | 609 | 601 |
| **Vip;Ai148** | Vip-IRES-Cre; Ai148(TIT2L-GC6f-ICL-tTA2) | 14 | 59 | 505 | 497 |
| **Pvalb;Ai162** | Pvalb-IRES-Cre; Ai162(TIT2L-GC6s-ICL-tTA2) | 7 | 39 | 299 | 299 |

## Results

### Characteristics of saccades in head-fixed mice during diverse visual stimuli

To better understand how mice in head-fixed contexts make saccadic eye movements, we extracted the eye position from eye tracking videos recorded during simultaneous stimulus presentation on a monitor placed contralateral to the recorded hemisphere of the mouse (de Vries et al., 2020). Specifically, we used DeepLabCut (Mathis et al., 2018) to fit ellipses to the eye and pupil in each frame of the videos, from which we calculated the pupil position in degrees (Fig. 1a; Methods). From this, we identified the occurrence of saccades by finding consecutive frames where the eye speed exceeded a threshold (Methods).

**Figure 1:**
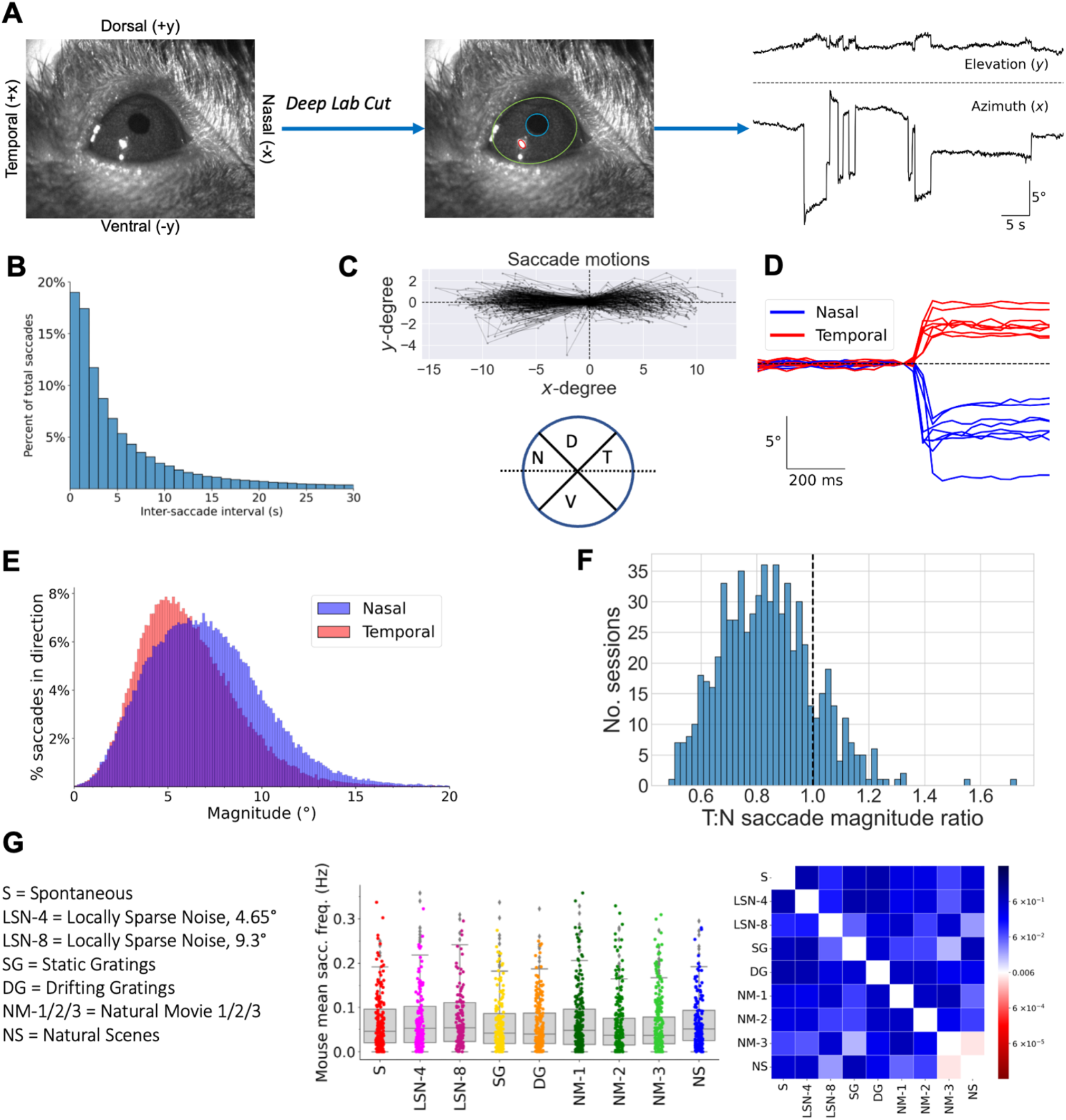
Saccade behavior is not influenced by visual stimulus. **(A)** (Left) Example frame from eye video annotated with axes of eye movement (x-axis nasal-temporal; y-axis dorsal-ventral). (Middle) Example ellipse fits on eyeball (green), pupil (blue), and glare spot (red) from DeepLabCut model (Methods). (Right) Example azimuth (horizontal) and elevation (vertical) traces from an experiment session after pupil extraction (Methods). Top trace is the elevation (y-degree, vertical), bottom trace is the azimuth (x-degree, horizontal), dashed line indicates 0°. Upwards is positive and downwards is negative using the same signs as in the left subplot. Times of abrupt eye position change correspond to saccades. **(B)** Inter-saccade intervals (s), defined as the time elapsed before the preceding saccade, for all saccades. Out of n = 202,156 total saccades, 33.5% saccades occurred at most two seconds after a previous saccade. **(C)** (Top) Example saccade motions from one experiment session, where each saccade has been aligned to start at the origin. Each saccade is a separate line, and points on the line indicate eye position at frames during the saccade. Note that saccades are made preferentially in the horizontal direction. (Bottom) Direction classification of saccades. Each sector of the circle is 90 degrees; dashed black line indicates horizontal axis. Middle intersection point denotes eye position alignment of the start of saccade. T, D, N, and V denote temporal, dorsal, nasal, and ventral, respectively. **(D)** Example nasal and temporal saccades from one experiment session. Dashed line represents 0°, and traces are centered by subtracting the position at the time of saccade onset. Only azimuthal (horizontal) traces are shown. **(E)** Distributions of saccade magnitude for nasal and temporal saccades. Nasal saccades (n = 104,121): 7.10° ± 3.19° (mean ± s.d.). Temporal saccades (n = 91,037): 6.14° ± 2.78°. Grouped together, all horizontal saccades (n = 195,158) had magnitude 6.65° ± 3.05°. Density normalization accounts only for saccades in the given direction; i.e., y-axis is normalized so both nasal and temporal area curves have unit area. **(F)** Ratio of mean temporal saccade magnitude to mean nasal saccade magnitude in each individual session. Dashed vertical line at 1, indicating an equal magnitude of nasal and temporal saccades. **(G)** (Left) Visual stimuli abbreviations used in this figure and throughout the paper. (Middle) Saccade frequency for different visual stimuli. Each data point is a mouse’s average saccade frequency for a particular visual stimulus (Methods). Box line corresponds to 25-75 percentile, line corresponds to median (50%), and error bars at 10% and 90%. (Right) Probability heatmap matrix comparing mouse mean saccade frequencies across different stimulus types using two-sample Kolmogorov-Smirnov (KS) test. Heatmap is centered at Bonferroni-corrected significance value (p = 0.05/8 ≈ 6e-3).

Across 818 one-hour experimental sessions, we identified a total of 202,156 saccades. The distribution of time elapsed between successive saccades had a long right tail, consistent with previous results that suggest saccades often occur in bursts separated by longer periods of fixation (Fig. 1b) (Samonds et al., 2018). Specifically, roughly one third (33.4%, n = 67,411) of all saccades were preceded by another saccade within two seconds.

We classified the direction of a saccade based on the direction of eye movement (Fig. 1c,d). Consistent with other studies of head fixed mice (Samonds et al., 2018; Meyer et al., 2020; Miura & Scanziani, 2022), we found that saccades were made almost exclusively along the horizontal axis. Specifically, 96.5% (n = 195,158) of all saccades ended within 45° from horizontal, with slightly more made in the nasal direction (51.5% [n = 104 121] nasal, 45.0% [n = 91,037] temporal). Also consistent with previous studies in both freely-moving (Meyer et al., 2020) and head-fixed (Sakatani & Isa, 2007; Itokazu et al., 2018; Miura & Scanziani, 2022) mice, we observed a systematic asymmetry in the magnitudes of nasal (7.10° ± 3.19°, mean ± s.d.) and temporal (6.14° ± 2.78°) saccades (Fig. 1e; mean ± s.d.; p = 0.0, two-sample Kolmogorov-Smirnov (KS) test). This discrepancy in magnitude was also found within individual sessions: the ratio of mean temporal saccade magnitude to mean nasal saccade magnitude within individual sessions was 0.84 ± 0.17, suggesting further that nasal saccades are slightly larger in magnitude than temporal saccades (Fig. 1f).

Given that the Visual Coding dataset records neural responses to a variety of visual stimuli— including drifting gratings (DG), static gratings (SG), locally sparse noise (LSN), natural scenes (NS), natural movies (NM-1, NM-2, and NM-3), and spontaneous activity (S)—we considered whether mice might make more saccades during certain visual stimuli by comparing the saccade frequencies during each of the visual stimuli (Fig. 1g). The distributions were heavily overlapping, indicating the stimulus did not heavily influence the saccade frequency. The only pairing of statistically-significant distributions was between natural scenes (NS) and natural movie 3 (NM-3), suggesting that mice may make slightly more eye movements during NS compared to NM-3. Given that there are no significant differences between in saccade distributions during NS and NM-1 or NM-2, however, it seems unlikely that this a meaningfully robust result.

### Neurons in visual cortex respond to saccades

To identify saccade-responsive neurons, we examined the dF/F traces of neurons surrounding each saccade. Our analysis included 32,442 neurons imaged in one or more imaging sessions (Methods). To compare our data with previous studies (Itokazu et al., 2018; Miura & Scanziani, 2022), we performed analogous tests to identify neurons with significantly modulated activity following saccades, directly comparing the dF/F activity following each saccade with activity preceding each saccade (Methods). Using these tests, we found that 20,629 neurons (63.6%) had significantly modulated activity following saccades, an estimate that is consistent with this previous work.

However, due to the high variability of neural activity over time, it is possible for activity to have statistically-significant differences in pre-saccade vs. post-saccade time intervals, but to fail to achieve significance when compared to randomly located moments in time. Indeed, from visual inspection it was clear that many of these modulated neurons did not exhibit clear and robust enhanced (or suppressed) responses to saccades (Fig. 2a). Thus, to identify neurons with robust saccade responses, we used a stricter bootstrapping procedure to estimate the null distribution of dF/F changes at randomly sampled (with replacement) time points throughout the entire experiment (Methods). This procedure identified saccade-responsive (SR) neurons across all cortical areas and layers that exhibited robust responses to eye movements in a variety of manners. Overall, 9.5% (n = 3,071) of analyzed neurons were SR. The saccade responses of SR neurons fell primarily into four distinct categories. First, 29.7% (n = 911) of SR neurons exhibited a strong enhanced (positive) response to all saccades, regardless of their direction (Fig. 2b). Second, 12.2% (n = 375) of SR neurons exhibited a strong suppressed (negative) response to all saccades (Fig. 2c). Finally, other SR neurons exhibited an enhanced response to saccades made along either the nasal or temporal direction, but no (or a slightly suppressed) response to saccades in the opposite direction (Fig. 2d,e; 10.3% [n = 318] nasal; 47.8% [n = 1,467] temporal). We refer to such neurons as direction-selective (DS) and define the direction—either nasal or temporal—that maximizes their response to be their preferred direction.

**Figure 2:**
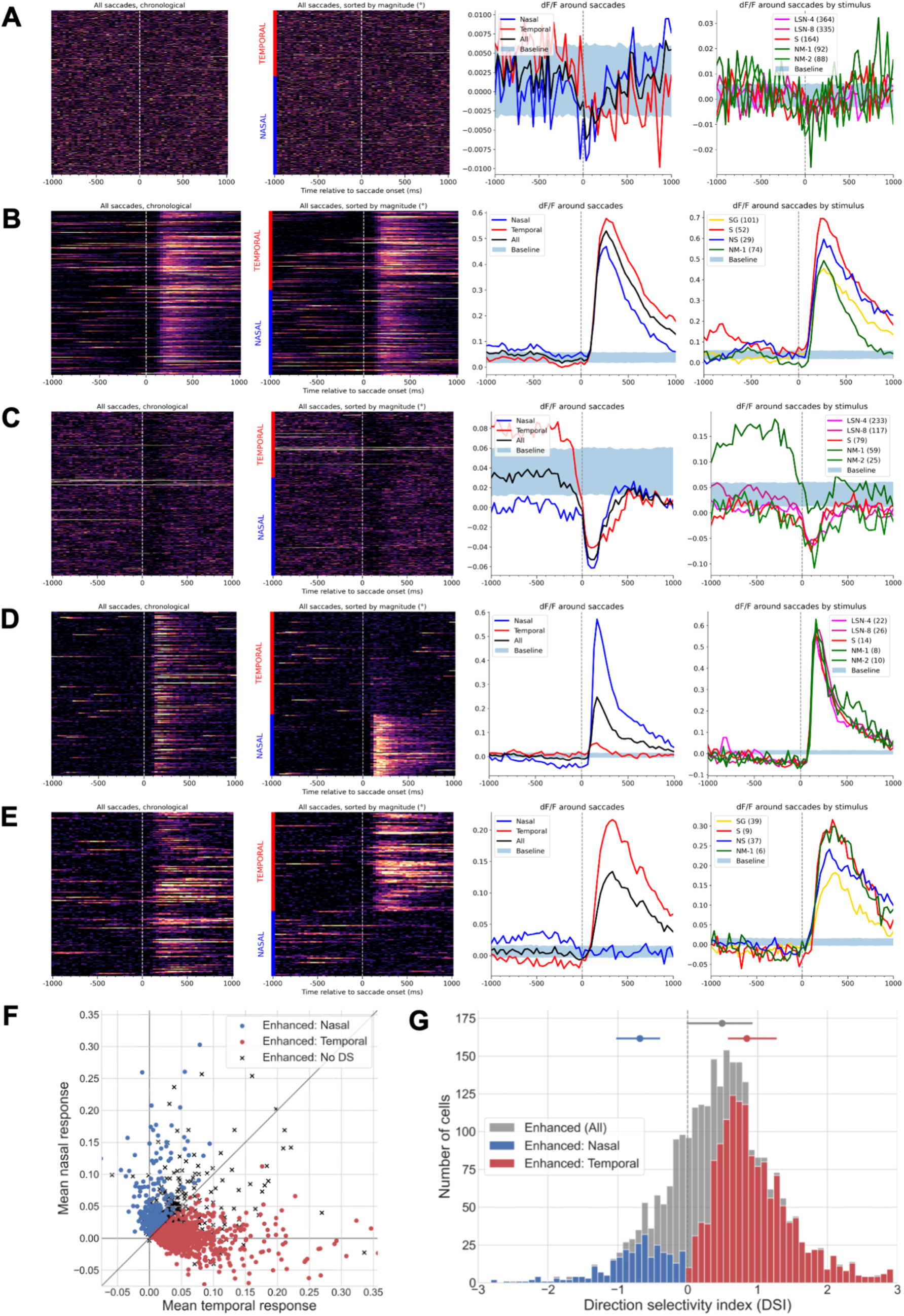
Diverse responses of neurons in visual cortex to saccades. **(A)** Example neuron with a significant activity modulation (Methods; p = 0.0024, Wilcoxon signed-rank). Neuron 669928467 (Tlx3;Ai148, VISl, L5, session 658854486). (Left) Each row is a dF/F trace centered around a saccade, where saccades are ordered in chronological order (top corresponds to start of experiment) and time zero corresponds to the start of each saccade. (Second from left) All temporal and nasal saccades (indicated by red and blue bars), where each group is sorted by magnitude (largest magnitude on top). (Third from left) Mean dF/F of neuron around saccades. Nasal (blue) and temporal (red) traces are mean dF/F around nasal and temporal saccades, respectively; all (black) corresponds to dF/F average around all saccades. Baseline is a N=1000 bootstrapped distribution where each trace is the mean of random time points within the experiment session (the number of random samples equals the number of saccades). (Right) dF/F average around preferred saccades by stimulus type. Baseline is the distribution from the previous subplot; number in legend is number of preferred saccades during each stimulus. **(B)** Example saccade-responsive (SR) neuron with an enhanced response to saccades, irrespective of direction. Neuron 517476630 (Cux2;Ai93, VISal L2/3, session 506156402). **(C)** Example SR neuron with a suppressed response to saccades, irrespective of direction. Neuron 670074250 (Cux2;Ai93 VISrl L2/3, session 662960692). **(D)** Example SR, direction-selective (DS) neuron preferring nasal saccades. Neuron 662076627 (Ntsr1;Ai148, VISp L6, session 606227591). **(E)** Example SR, DS neuron preferring temporal saccades. Neuron 662209107 (Ntsr1;Ai148, VISp L6, session 636889229). **(F)** Scatter plot of saccade-responsive enhanced neurons showing their mean temporal saccade response (x-axis) and mean nasal saccade response (y-axis). Blue dot indicates neuron prefers nasal saccades, red dot indicate neuron prefers temporal saccades, black x indicates neuron is not direction-selective. Mean nasal/temporal response for nasal neurons: 0.0528/0.0074, temporal neurons: 0.0015/0.0490; non-direction-selective neurons: 0.0249/0.0302. **(G)** Direction selectivity index histogram for saccade-responsive enhanced neurons. Gray: all saccade-responsive enhanced neurons (n = 2,696), blue: neurons that prefer nasal saccades (n = 318), red: neurons that prefer temporal saccades (n = 1,467). Bar is middle 50% (all: [-0.01, 0.93], nasal: [-1.02, −0.39], temporal: [0.58, 1.27]); dot is median (all: 0.49, nasal: −0.68, temporal: 0.84).

In our analysis, SR neurons were classified as DS if they met the following two criteria: (i) have an enhanced response to saccades, and (ii) have a statistically significant difference in response to nasal and temporal saccades (Methods; p < 0.05, Wilcoxon rank-sum test). A DS neuron was further classified as preferring temporal saccades if *T* > *N*, where *T* was its mean response to temporal saccades and *N* was the mean response to nasal saccades, or as preferring nasal saccades if *N* > *T* (Fig. 2f). To quantify direction preference, we calculated a direction selectivity index (DSI) metric for DS neurons, defined as (*T* – *N*)/(*T* + *N)* (Fig. 2g). Note that given *T* and *N* can take on negative values our DSI metric is not necessarily constrained between −1 and + 1; furthermore, a positive DSI indicates a temporal saccade preference, and a negative DSI indicates a nasal saccade preference. The mean DSI of temporal-saccade-preferring neurons was 1.06 ± 1.56, and that of nasal-saccade-preferring neurons was −0.88 ± 1.03 (mean ± s.d.). Surprisingly, we observed a strong bias for DS neurons to prefer temporal saccades over nasal saccades (temporal, n = 1,467 neurons; nasal, n = 318 neurons), a result that has not been discussed in prior saccade literature. Applying the DSI computation to all enhanced SR neurons, we found the mean to be 0.50 ± 1.41, suggesting that even non-DS SR-enhanced neurons responded more strongly on average to temporal saccades than nasal saccades. This bias was not due to a discrepancy in the quantity of nasal and temporal eye movements within sessions, as 47.6% ± 12.9% (mean ± s.d.) saccades in each experiment were nasal, and 47.2% ± 11.5% were temporal. This even split between nasal and temporal saccades suggests the bias toward neurons preferring temporal saccades is a result of neural dynamics rather than behavioral tendencies.

How individual SR neurons vary their responses during visual stimulation might inform alternative models for how cortex integrates saccades in the presence of visual stimuli. Generally, SR neurons exhibited strong responses to saccades independent of visual stimulus, but the degree of that response for particular neurons was sometimes enhanced or suppressed by different visual stimuli. To quantify this variability, we measured the saccadic response during particular visual stimuli relative to the size of the saccadic response during a mean-luminance gray screen (the “spontaneous” stimulus). We only considered experimental sessions where at least 8 saccades were made during the spontaneous stimulus (Methods). Furthermore, for DS neurons, we investigated only those saccades made in the neuron’s preferred direction. We first compared the mean saccade response to saccades made during the spontaneous stimulus to the mean saccade response during the other visual stimuli (Fig. 3a; Wilcoxon signed-rank test). We found no significant difference in saccade responses during artificial stimuli (LSN-4, LSN-8, DG, SG) compared to responses to saccades made during the spontaneous stimulus. However, we observed significant differences in the saccade responses between spontaneous and naturalistic stimuli (NM-1, NM-2, NM-3, and NS), suggesting more substantial heterogeneity in saccade responses during these visual stimuli. To better understand how these responses varied by stimulus, we compared the ratio of the mean response to saccades made during NM-1 to the mean response to saccades during spontaneous (Fig. 3b; Fig 3., Supplement 1). While there was considerable variation in these ratios, the median ratio for every stimulus was less than 1 (median of 0.87, 0.86, 0.86, 0.82, 0.95, 0.82, 0.92, 0.91 for LSN-4, LSN-8, SG, DG, NM-1, NM-2, NM-3, and NS, respectively), suggestive of a diminished response to saccades made during visual stimuli when compared to the response to saccades made during a gray screen. This indicates that while there is great heterogeneity in saccade responses of SR neurons, these responses are marginally suppressed during naturalistic—but not artificial—stimuli. Recalling how we define a saccadic response—the difference in neural activity around the saccade—the observed diminished saccade response during natural stimuli may in part be caused by elevated cortical activity and increased responsiveness of neurons during natural versus artificial stimuli (de Vries et al., 2020).

**Figure 3:**
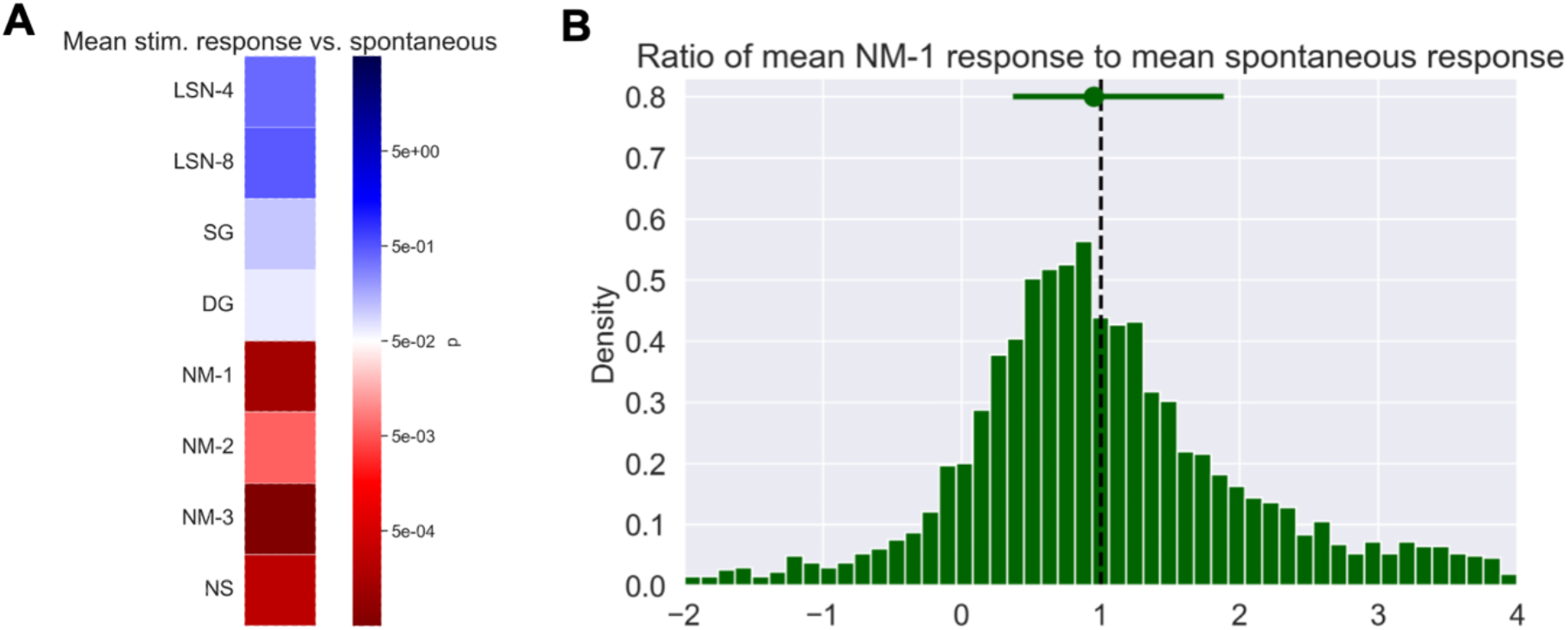
Saccade responses are suppressed by naturalistic but not artificial stimuli. **(A)** Comparison of mean response to saccades made during spontaneous visual stimulus and other visual stimuli (Methods). Significance value is p = 0.05 using Wilcoxon rank-sum test. **(B)** Ratio of mean response to preferred saccades made during NM-1 to mean response of preferred saccades made during spontaneous for each saccade-responsive neuron. Vertical black dashed line at 1 indicates equal response. Green dot is median ratio, bars extend to 25% and 75%. For additional stimuli, refer to Figure 3, Supplement 1.

### Differences in saccade response across transgenic lines and cortical areas

The scale of the Brain Observatory dataset enables us to compare saccade responses across neurons in different visual areas, transgenic lines, and cortical layers (Table 1). Saccade-responsive neurons were found across all visual areas, transgenically-defined cell types, and cortical layers— though not at the same frequency across these dimensions (Fig. 4a). Sst;Ai148 neurons exhibited the highest overall SR frequency, at 22.5% (135 out of 601), whereas Slc17a7;Ai94 neurons exhibited the lowest overall SR frequency, at 2.6% (21 out of 817). We observed higher proportions of both SR and SR-suppressed neurons in TIGRE 2.0 (Ai148 or Ai162; (Daigle et al., 2018)) reporters than the TIGRE 1.0 (Ai93 or Ai94; (Madisen et al., 2015)) reporters (Fig. 4b). As there is no Cre diver that was used with both TIGRE 1.0 and TIGRE 2.0 constructs, however, we are unable to determine whether this difference is because TIGRE 2.0 was used specifically for inhibitory neurons and excitatory neurons in deep layers, or if this is the result of an off-target effect of the over-expression of either tTA or GCaMP using the TIGRE 2.0 reporter. Additionally, DS SR neurons exhibited a noticeably higher preference for temporal saccades over nasal saccades, and this observation was consistent across all lines.

**Figure 4:**
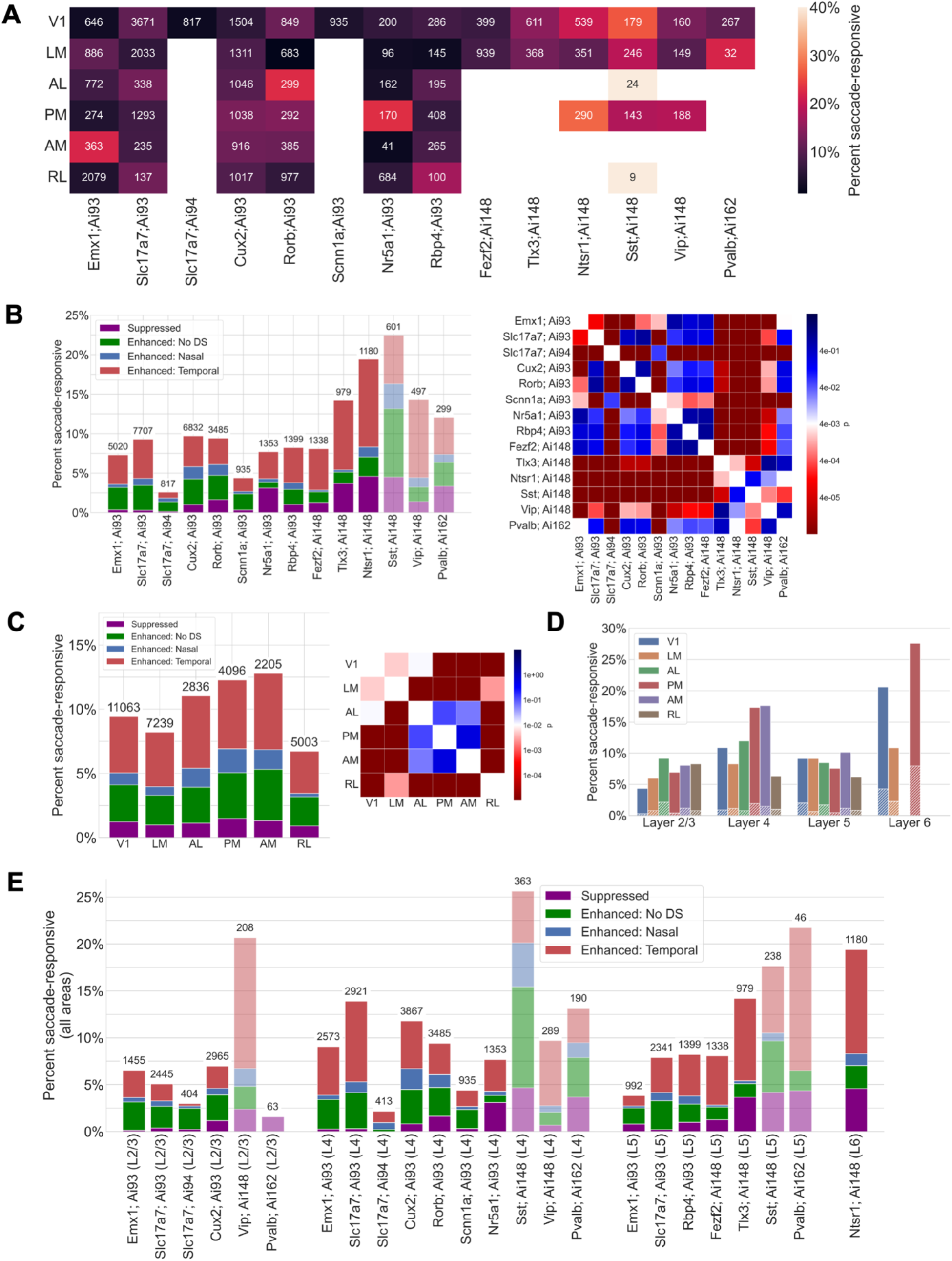
Differences in saccade responses across transgenic lines and visual areas. **(A)** Heatmap of percent of saccade-responsive (SR) neurons by transgenic line and visual area, relative to total number of imaged neurons, which is given above each bar. **(B)** Left: Percent of SR neurons by transgenic line, broken down by type of response. “Enhanced: No DS” means the neuron has an enhanced response to saccades but does not prefer any one direction (Methods). Number above each bar is the total number of imaged neurons of the corresponding line. Low opacity bars indicate inhibitory neurons. Right: Chi-squared test across pairings of transgenic lines (using a 2×2 contingency matrix containing number of SR and non-SR neurons for each line). **(C)** Left: Percent of SR neurons by visual area. Number above each bar is the total number of imaged neurons in the corresponding area. Right: Heatmap; same as in (B). **(D)** Percent SR neurons by cortical layer and visual area. Hatched marks indicate neurons with a suppressed saccade response. **(E)** Percent SR neurons by transgenic line and cortical layer, across all visual areas. Low opacity bars indicate inhibitory neurons, and numbers above bars indicate the total number of neurons imaged.

We also observed high percentages of saccade responsive neurons in anterolateral (AL), anteromedial (AM), and posteromedial (PM) areas (Fig. 4c), and these rates were consistent across these three areas and statistically different from other areas (chi-squared test, p < 0.01). This higher rate of SR neurons in these dorsal areas is consistent with dorsal areas representing motion information or egocentric reference frames (Andermann et al., 2011; Marshel et al., 2011; Wang et al., 2012). Rostrolateral (RL) neurons exhibited the lowest percentage of SR neurons, at 6.7% (336 out of 5,003). However, this discrepancy to the other dorsal areas above (that exhibited greater proportions of saccade responsive neurons) could reflect the challenges of accurate retinotopic targeting of RL neurons (de Vries et al., 2020; Siegle et al., 2021).

Next, we investigated the distribution of SR neurons by layer. Previous work has shown that saccade-induced neural responses increase in excitability and discriminability as a function of cortical depth (Miura & Scanziani, 2022). We similarly find variations in the percentage of SR neurons by layer, with high percentages for neurons in layer 6 (Fig. 4d). In this deep layer, Ntsr1;Ai148 neurons imaged in V1, LM, and PM exhibited the highest overall frequencies of saccade responsivity. Interestingly, a higher proportion of SR layer 6 neurons exhibited a suppressed response relative to more superficial layers. We also found a high frequency of SR neurons in layer 4 PM and AM neurons. In aggregate, only a small fraction of superficial layer 2/3 neurons was SR, and this was consistent across all visual areas.

Finally, as some transgenic lines were imaged in different layers, we examined this layer-dependent breakdown across different transgenic lines (Fig. 4e). Broadly, excitatory neurons in deeper layers (e.g., Tlx3;Ai148 in layer 5 and Ntsr1;Ai148 in layer 6) tended to be more saccade-responsive, but inhibitory neurons did not exhibit this same bias toward deeper layers. Specifically, across all visual areas, 20.7% (40 out of 208) Vip;Ai148 inhibitory layer 2/3 neurons were SR, 25.6% (93 out of 363) Sst;Ai148 inhibitory layer 4 neurons were SR, and 21.7% (10 out of 46) Pvalb;Ai162 inhibitory layer 5 neurons were SR—these were the highest rates across any population in all layers. In addition to these inhibitory populations showing higher proportions of suppressed saccade responses, they also tended to prefer specific layers, with Vip;Ai148 preferring layers 2/3, Sst;Ai148 preferring layer 4, and Pvalb;Ai162 preferring layer 5. This layer preference was also prevalent in some excitatory populations, with the most notable difference occurring in layer 5. Here, 14.2% (139 out of 979) Tlx3;Ai148 neurons were SR, while only 8.1% (108 out of 1338) Fezf2;Ai148 were SR. Tlx3;Ai148 neurons are cortico-cortical projecting neurons (Gerfen et al., 2013) while Fezf2;Ai148 neurons are corticofugal (Guo et al., 2013), suggesting that the saccadic activity is differentially represented in these distinct pathways. These cell type dependencies were also consistent when examining primary visual cortex specifically (Fig. 4, Supplement 1), with a minor difference being V1 layer 2/3 excitatory cells were slightly less likely to be saccade-responsive when compared to rates across all visual areas.

### Saccade responsive neurons have similar visual responses as non-saccade responsive neurons

Given that visual cortex receives both bottom-up visual input from the eyes and top-down feedback from other areas, a key question about saccade responses is whether they are driven by the change in visual features due to the eye movement, motor signals, or a combination of both. To consider this, we compared the visual response properties of SR neurons with those of non-SR neurons. If saccade responses are primarily visually-driven, we would expect SR neurons to exhibit visual tuning properties that are distinct from those of non-SR neurons.

Previous work had used a cluster analysis of the response reliabilities of neurons to four of the stimuli (DG; SG; NS; and NM, combining together NM-1, NM-2, and NM-3) to identify functional response classes with this dataset (de Vries et al., 2020). This analysis categorized neurons based on the stimuli to which they responded reliably. Broadly, neurons fall into 9 of the possible 16 combinations of stimuli, including a “None” class in which neurons do not reliably respond to any of these visual stimuli. We examined how SR neurons were distributed across these functional classes and found that each of the nine classes was composed of roughly 10% SR neurons (Fig. 5a). This observation that SR neurons are not biased toward a single functional response class suggests that SR neurons do not exhibit a particular visual response feature that is driven by saccades. We used a chi-squared test with a 2×2 contingency table containing the number of SR and non-SR neurons in each cluster to compare these rates across classes, finding no significant differences (*p* > 0.05, Bonferroni correction), except in one case between “None” and “NM” classes, suggesting neurons responsive to no visual stimuli may be slightly more likely to respond to saccades than neurons responsive to only natural movies.

**Figure 5:**
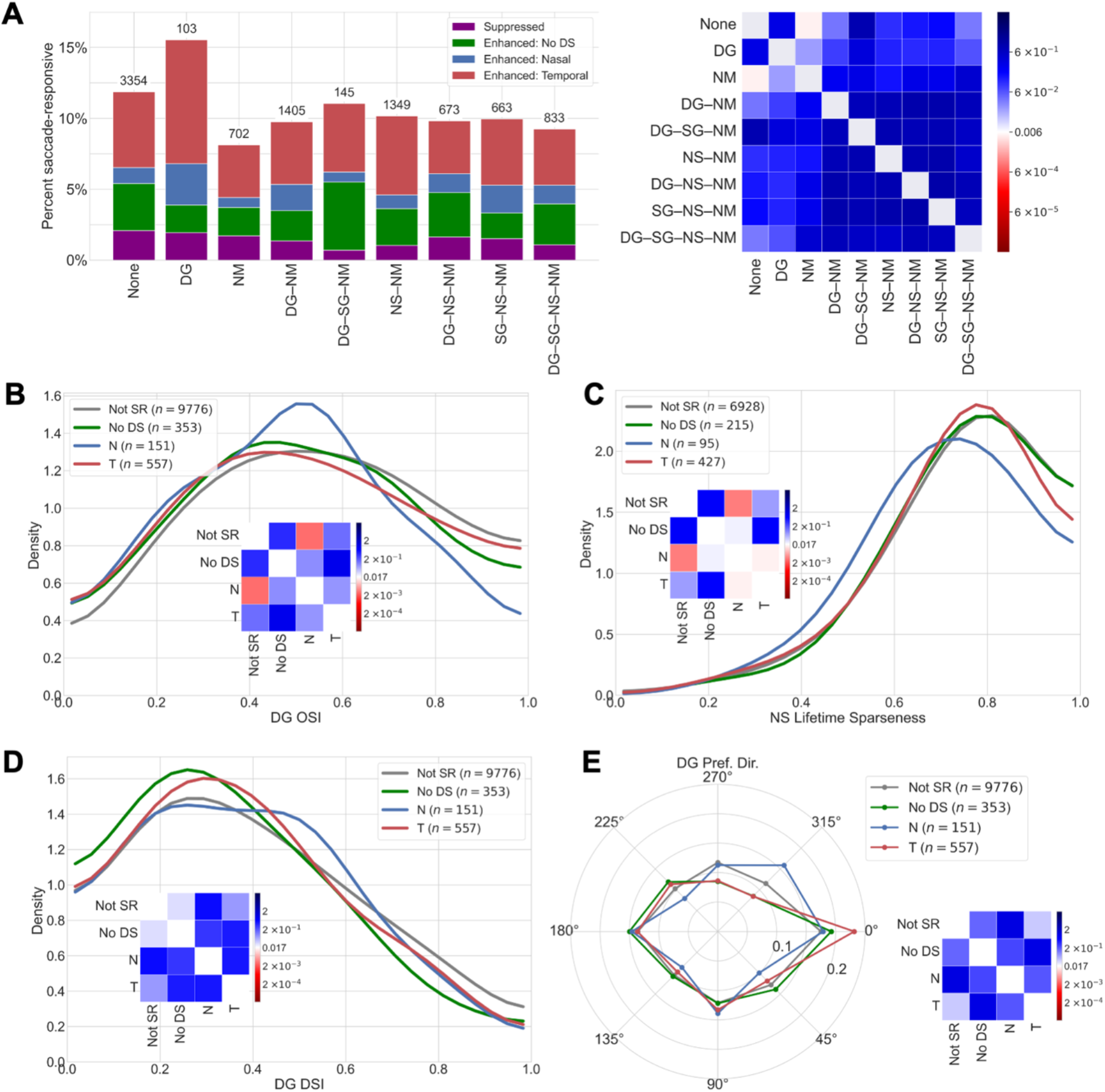
Saccade-responsive neurons have similar visual responses to non-saccade-responsive neurons. **(A)** (Left) Percent of SR neurons by visual response class, indicating which set of visual stimuli elicit a response. Number above each bar is the total number of imaged neurons within the given cluster. (Right) Chi-squared test across pairings of transgenic lines (contingency matrix containing number of SR and non-SR neurons for each cluster; p = 0.05, Bonferroni corrected for multiple comparisons). **(B)** Orientation selectivity index for neurons that respond to drifting gratings. Curve is smoothed using a Gaussian kernel. Inset: Heatmap shows KS test between different distributions (p = 0.05, Bonferroni corrected for multiple comparisons). **(C)** Analogous plot to (B), showing lifetime sparseness for neurons responsive to natural scenes. **(D)** Analogous plot to (B), showing direction selectivity index plot for neurons responsive to drifting gratings. **(E)** Distribution of preferred direction for neurons responsive to drifting gratings.

Given we observe no clear preference for functional visual response class among SR neurons, we sought to identify any differences in broader tuning metrics for SR neurons to different visual stimuli. We first investigated neurons responsive to drifting gratings, comparing the orientation selectivity index (DG OSI) metric measuring how narrowly tuned a neuron is for oriented drifting gratings (1 = narrowly tuned, 0 = broadly tuned; Methods). SR and non-SR neurons had similar OSI distributions (Fig. 5b); the only significantly different comparison was between the OSI of non-SR neurons and that of nasally-selective SR neurons, suggesting nasally-selective SR neurons may to a lesser extent discriminate the orientation of the DG stimulus. However, this difference may be caused in part by the small number of nasally-selective SR neurons (n = 95) relative to other categories, and overall reflects negligible differences in orientation selectivity among SR and non-SR neurons. Next, we investigated neurons responsive to natural scenes (118 different natural scene visual images shown to mice every 250 ms), comparing the lifetime sparseness metric neuron capturing how selective a neuron is to the images shown (1 = responsive only to a single image, 0 = broadly responsive to all images) (Vinje & Gallant, 2000; de Vries et al., 2020). If SR neurons had significantly lower lifetime sparseness values compared to non-SR neurons, then this may suggest they are more broadly responsive to abrupt changes in visual stimulus (such as those induced by a rapid eye movement). On the contrary, we found all lifetime sparseness distributions to be rightward skewed and heavily overlapping among SR and non-SR neurons (Fig. 5c). However, nasally-selective SR neurons exhibited a slightly and statistically-significantly leftward-skewed lifetime sparseness distribution, suggesting these neurons may respond more broadly to visual stimuli, though again this difference may be due to small sample size. All other distributions of non-SR neurons, temporally-selective SR neurons, and non-DS SR neurons overlapped, suggesting these latter types of SR neurons exhibited similar responses to visual scenes to those of non-SR neurons.

We finally considered the possibility that SR neurons were responding to the saccade-induced translation of the visual field. Specifically, if saccade responses are driven by visual field motion, we would expect SR neurons to respond preferentially to visual motion along the horizontal axis, as this is the primary axis of saccadic eye movements. Furthermore, we would expect DS SR neurons to respond to visual motion in the opposite direction as their preferred saccade direction, since this is the direction of visual field motion when the eye moves in the preferred saccade direction. To identify whether this was the case, we analyzed drifting grating tuning metrics, first comparing the direction selectivity index (DG DSI), a metric comparing the neuron’s response to its preferred and non-preferred directions of motion for the drifting gratings stimulus (Fig. 5d; Methods). We found no significant difference between the distributions of SR neurons and non-SR neurons, suggesting SR neurons are not differentially selective to specific directions of motion (two-sample KS test, *p* > 0.05, Bonferroni correction). Finally, we compared the preferred drifting grating direction of motion (the direction that maximizes the neural response) across neurons (de Vries et al., 2020). Again, we found no significant differences in these distributions between SR and non-SR neurons, suggesting SR neurons do not differentially prefer motion along the horizontal axis (Fig. 5e). These two results taken together indicate there are no notable differences in visual motion responses of SR neurons relative to those of non-SR neurons, suggesting the saccade-induced visual field motion does not primarily drive the responses of SR neurons.

In summary, this analysis of visual responses finds that SR neurons do not exhibit distinct response properties from the overall neural population, suggesting that saccade responses are not primarily visually-driven.

## Discussion

In this study, we examined saccadic eye movements and visual cortical neural responses in head-fixed mice. These mice tended to make spontaneous saccades, with a frequency independent of the visual stimulus, occurring frequently in successive bursts, and preferentially along the horizontal direction. Saccade-responsive (SR) neurons were found across all cortical areas, layers, and transgenically-defined cell types that were sampled in this dataset, though with differing prevalence across these dimensions. Previous work has found the dynamics of SR neurons are similar in both freely moving and head-fixed mice (Miura & Scanziani, 2022), and this suggests the distributions we have found of SR neurons across cell type, cortical layer, and visual area should generalize to freely-moving mice. Overall, our analysis reveals novel insights about the distribution of SR neurons and lack of distinct visual response properties of these neurons across the visual cortex.

### Neural saccade response distributions across transgenic line, cortical layer, and visual area

SR neurons were found in all visual areas and all the of the cell types that were surveyed in this dataset, however there were notable differences in the prevalence of SR neurons. First, SR neurons occurred more frequently in dorsal visual areas AL/PM/AM. This is consistent with egocentric representations and spatial processing in dorsal visual areas, but future work examining saccade responses in more ventral visual areas is needed to determine if there is a clear difference in rates of saccade-responsive neurons in these two pathways. Second, inhibitory neural cell types (Sst;Ai148, Vip;Ai148, Pvalb;Ai162) exhibited consistently high SR rates across visual areas (Taniguchi et al., 2011). Past work has shown that the inhibitory microcircuit mediates both bottom-up and top-down effects on cortical representation (Zhang et al., 2014; Dipoppa et al., 2018), and future work is needed to fully understand how these inhibitory interneurons integrate saccade signals with other sensory and behavioral signals.

Finally, there was a bias for SR neurons to occur more frequently in deeper cortical layers rather than supragranular layers, consistent with findings in a recent electrophysiological study (Miura & Scanziani, 2022). Although aggregating all visual areas revealed many layer 4 excitatory cell types that responded to saccades (Fig. 4e), analysis in V1 specifically revealed lower SR rates in layer 4 (with the exception of Slc17;Ai93) and a prevalent SR bias toward cell types in layers 5 and 6 (Fig. 4, Supplement 1). These findings are consistent with anatomical inputs from LP to visual cortex, finding LP targets layer 5 in V1 and layer 4 in higher visual areas (Herkenham, 1980). We also found relatively high SR rates in Tlx3;Ai148 neurons in V1, which are cortico-cortical projecting layer 5 excitatory neurons that have been shown to both provide direct feedforward input to and receive feedback from higher visual areas, in addition to superior colliculus (Gerfen et al., 2013; Kim et al., 2015). This high rate, in contrast to the corticofugal Fezf2;Ai148 layer 5 excitatory neurons (Guo et al., 2013), may implicate their importance in modulating levels of feedforward inputs and suggests that saccadic activity is differentially represented in these distinct pathways. Additionally, the Ntsr1;Ai148 transgenic line, labeling the majority of excitatory layer 6 neurons (Gong et al., 2007), had the highest SR rate of any excitatory neural cell type. Given that a distinct subpopulation of these Ntsr1;Ai148 neurons have apical dendrites extending to L1 (Olsen et al., 2012), the high SR rate we observed in this population is consistent with the notion that LP relays saccade signals to visual cortex through projections to superficial cortical layers (Roth et al., 2016; Miura & Scanziani, 2022). Moreover, given that these L6 neurons project both to superficial layers and to primary sensory thalamic nuclei (Olsen et al., 2012), they may also play a role in modulating visual representations through both intracortical and thalamocortical pathways.

Nonetheless, we emphasize the anatomical reality that many cortical neurons have layer-spanning dendrites, and it would be incorrect to attribute the saccade-modulated activity of the neuron to the layer of its soma. Thus, future research is needed to localize precisely where saccade signals enter cortex and leverage these insights to elucidate the role of bidirectional corticothalamic feedback in regulating sensory processing during saccades.

### Saccade-responsive neurons biased toward preferring temporal saccades

Saccade-responsive neurons with enhanced, direction-selective responses overwhelmingly favored saccades made in the temporal direction, and this bias was consistent across transgenic lines, cortical layers, and visual areas. Though other asymmetries in nasal and temporal saccade magnitudes (Meyer et al., 2020) and velocities (Maruta et al., 2006) have been previously documented, our finding of visual cortical neurons disproportionately favoring temporal saccades has not been documented and raises interesting questions about neural responses in mice. We found there was no clear preference for mice to make saccades in either the nasal or temporal direction; saccades in these two directions were made at the approximately same rates in individual sessions. Furthermore, our analysis of visual tuning curves suggests these DS neurons do not differentially prefer horizontal visual motion in any specific direction, ruling out the explanation of this effect being due to a visual response bias toward horizontal motion.

One possible explanation, supported by our observation that the neural response was stronger with temporal saccades in all enhanced SR neurons (as opposed to only direction-selective neurons; Fig. 2g), involves differences in neural processing of monocular versus binocular vision. Mice have a central 40° binocular field of view, but the majority of their visual field is monocular, with retinal ganglion cells throughout the entire retina projecting to the contralateral hemisphere (Seabrook et al., 2017). Under this explanation, the bias toward preferring temporal saccades may be related to previously-documented sensitivity to binocular disparity that is widespread across mouse visual cortex (La Chioma et al., 2019). Another possibility is that because a fraction of nasal saccades in freely-moving mice may be compensatory for optic flow in self-motion, it may be that the visual system has evolved to differentially enhance the response of saccade responses in the temporal direction.

### Visual versus motor responses

Saccade-responsive visual cortical neurons have previously been observed to respond to a mix of visual and motor saccade responses (Itokazu et al., 2018; Miura & Scanziani, 2022). If our SR neurons were consistently responsive to no visual stimuli (i.e., in the “None” class of visual response), then this may suggest the saccade response is largely non-visual, as the neurons exhibit little stimulus-driven visual response. On the other extreme, SR neurons with visual response properties deviating substantially from non-SR neurons, along with differences between SR-nasal and SR-temporal neurons (such as a different direction selectivity or preferred direction for drifting gratings), would implicate a largely visually-driven saccade response.

However, we observed neither of these extremes (Fig. 5). First, SR neuron visual response properties overlapped heavily with those of non-SR neurons, indicating SR neuron visual response properties are not substantially different from what we would expect by taking a random subset of the neuronal population. This suggests that the saccade responses of the neurons we identified are not uniquely visually-driven, and conversely, neurons responsive to a particular visual stimulus are not differentially likely to respond to saccades. Second, SR neurons in our dataset tended to exhibit time-locked saccade responses during all visual stimuli—including the spontaneous stimulus (mean-luminance gray screen), during which there would be a negligible visual response induced by a change in the visual receptive field. These observations indicate that the saccade responses here are not visually driven, but instead reflect motor signals being propagated through the cortex.

We observed some minor heterogeneity, however in the magnitude of saccade-responses observed during different visual stimuli, particularly in SR responses to saccades made during the spontaneous stimulus and those made during naturalistic stimuli (Fig. 3; natural movies and natural scenes). These latter stimuli have richer and more dynamic spatial statistics than other stimuli, driving greater overall cortical activity in both excitatory and inhibitory populations (de Vries et al., 2020). This suggests that visual signals do modulate neural saccade responses, and supports a model of linear integration of these signals as proposed in (Miura & Scanziani, 2022).

## Acknowledgements

Research funding was generously provided through the University of Washington Levinson Emerging Scholars Program while the first author was an undergraduate at the University of Washington. We would also like to thank Alan Degenhart for his helpful insights while the first author was a summer intern at the Allen Institute, Dan Millman for his statistical advice, and Forrest Collman for his helpful feedback on initial manuscripts. We wish to thank the Allen Institute founder, Paul G. Allen, for his vision, encouragement, and support.

## Author contributions

**CK**: Conceptualization, Methodology, Software, Formal Analysis, Writing – Original Draft, Writing – Review & Editing, Visualization. **PL**: Data Curation, Software. **MAB**: Methodology, Writing – Review & Editing, Supervision. **SEJdV**: Conceptualization, Methodology, Writing – Review & Editing, Supervision.

## Methods

### Dataset

We used the 2-photon calcium imaging recordings from the Allen Brain Observatory Visual Coding dataset (*Allen Brain Observatory -- 2-Photon Visual Coding*, 2016; de Vries et al., 2020). This open dataset consists of neural activity recorded using 2-photon calcium imaging of transgenically expressed GCaMP6, surveying activity from neurons found in 6 cortical areas, 4 cortical layers, and 14 transgenic Cre lines. Awake, head-fixed mice were imaged while viewing a diverse set of visual stimuli including drifting gratings, static gratings, natural scenes, natural movies, locally sparse noise, and a mean luminance gray screen (“spontaneous activity”). These stimuli were divided across three ~1-hour imaging sessions, and each field of view was imaged across three days to record these distinct sessions. A set of three sessions from a given field of view is an “experiment container.” Cell matching was performed to match identified ROIs across the sessions within an experiment container. Neural recordings were made from the left hemisphere while the mouse viewed a monitor to its right side. To track the mouse’s eye movements, the right side of the mouse was illuminated with an infrared LED and the eye was imaged using an Allied Vision camera (Mako G-032B with GigE interface, 30 FPS). All experimental procedures are detailed in (de Vries et al., 2020). Table 1 lists the transgenic lines in the datasets and the number of experiment sessions analyzed in this study for each line.

### Stimulus abbreviations

LSN-4 and LSN-8 refer to Locally Sparse Noise, 4° and 8°, respectively. SG refers Static Gratings, DG refers Drifting Gratings. NM-1, NM-2, NM-3 refer to Natural Movies One, Two and Three respectively, and NS refers to Natural Scenes. The stimulus parameters are detailed in (de Vries et al., 2020)

### Analysis

All analyses were performed using custom scripts written in Python 3.7, using NumPy (Harris et al., 2020), SciPy (Virtanen et al., 2020), Pandas (McKinney, 2010), Matplotlib (Hunter, 2007), and the AllenSDK.

### Eye video trace extraction

We trained a single, universal eye tracking model in DeepLabCut (Mathis et al., 2018), a ResNET-50 based network, to recognize up to 12 tracking points, each, around the perimeter of the eye, the pupil, and the corneal reflection, respectively. We used a published numerical routine to fit ellipses to each set of up to twelve tracking points (Hammel & Sullivan-Molina, 2020). For each ellipse, we reported the following parameters: (x, y) coordinates of the ellipse center, the (width, height) half-axis of the ellipse, and the rotation angle *ϕ* around the center. Fits were performed individually on each frame if there were at least 6 tracked points with a confidence of at least 0.8 as reported by the output of DeepLabCut. For frames where there were fewer than 6 tracked points, the ellipse parameters were set to not-a-number (NaN).

The training data set contained two sources of hand-annotated data. We uniformly sampled 40 (out of 1848) behavior movies at random and extracted 3 frames from each (for a total of 120 frames) via the *kmeans* option in DeepLabCut for annotation. On each frame, 8 points each were annotated, approximately equidistantly around the eye and pupil, respectively. The center of the corneal reflection was annotated with a single point. These labeled frames comprised data set (1). Next, we manually annotated 4150 frames with a rotated ellipse fitting the pupil and a point for the corneal reflection; the perimeter of the eye was not annotated. These labeled frames formed data set (2). Then, in order to obtain a training data set where twelve tracking points were annotated for each the eye, the pupil, and the corneal reflection, respectively, we bootstrapped the above training data as follows. We trained a model on (1) and used it to track the eye outlines on the data in (2). This yielded 4150 frames fully-annotated with ellipses (human annotation for pupil and corneal reflection, DeepLabCut annotation for the eye). Twelve points, in angular increments of 30 degrees were computed along each fitted ellipse for use as training data set (3). A new model was trained on this new data set (3); 100 frames were extracted from each of the 40 videos used to create (1), and the new model was used to fit 3 × 12 points to the 4000 frames obtained in this manner yielding data set (4). The data sets (3) and (4) were combined, and the final, universal eye tracking model was trained on the combined 8150 frames for 1,030,000 (DeepLabCut’s default maximum number of) iterations.

### Automated detection of saccades

Using the extraction pipeline detailed above, we computed the absolute movement speed (°/s) along for pupil azimuth and elevation using a discrete central difference approximation for the first derivative:

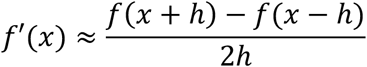

Then, we combined these two dimensions using the Pythagorean theorem to determine the absolute eye speed (°/s). Following a similar approach as previous studies (Ramirez & Aksay, 2021), we defined the eye speed outlier threshold as μ + 3σ or 10 °/s, whichever was higher (where statistics are computed in the entire trace during the experiment, ignoring NaN frames), and identified all of the frames during the experiment when the eye speed exceeded this threshold. Around each of these frames, we identified the frames where the speed exceeds μ + 1σ, resulting in a start and end frame for each saccade.

To remove saccades that were flagged incorrectly due noisy position traces, eye bulging, or other animal jitter, we computed the mean and standard deviation of the eye position trace in the 300 ms before saccade onset and 300 ms after saccade offset, for both azimuth and elevation. We ignored saccades that had any dropped data in these traces. We then computed two thresholds of 3 · *max*(σ_before_, σ_after_) along these two dimensions to quantify the amount of noise in the trace; a larger quantity indicated a more substantial amount of noise in the trace. We computed the absolute difference in eye position during the flagged saccade, and the saccade was only considered valid if this difference exceeded the threshold in at least one dimension.

We computed the direction of each saccade by comparing the eye position at the saccade offset to the position at the onset. If the angle fell in the 90° sector between 45° and 315° (i.e., between −*π*/4 and *π*/4), the direction was set as temporal (right); if the angle fell in the 90° sector between 135° and 225° (i.e., between 3*π*/4 and 5*π*/4), the direction was set as nasal (left).

### Saccade analysis

From each experimental session, we identified which visual stimulus was present during each saccade. Since each individual mouse was imaged in one to three sessions, we aggregated the saccade frequency per stimulus statistic across these multiple sessions by dividing the total number of saccades made during a stimulus by the total duration the mouse was shown the stimulus.

### Data inclusion criteria

To analyze the saccade responses, we combined the experiment sessions from each experiment container to use all saccades for each neuron. Saccades that were not preceded by at least two seconds of steady gaze (i.e., no saccades) were excluded. Experiment containers with less than 15 nasal saccades or less than 15 temporal saccades were excluded. Of thedataset containing 43,316 neurons, we analyzed data from 32,442 neurons across 205 mice and 818 sessions (Table 1).

### Saccade visual activity modulation

To compare our results to recent work done by (Miura & Scanziani, 2022), we performed two analogous tests to identify neurons with significantly modulated responses. The first test was a Wilcoxon signed-rank test comparing the mean dF/F after the saccade onset (frames 0 to 10 relative to the saccade onset, i.e., 0 to 333ms after saccade start) to the mean dF/F before the saccade (frames −20 to −10 relative to the saccade onset; 667ms to 333ms before saccade start). The second test was a Wilcoxon rank-sum comparing the mean dF/F after the saccade onset (same range) for saccades in nasal vs. temporal directions. The significance threshold on both tests was set as p = 0.05, and a neuron was considered to have significantly modulated visual activity if there was significance in at least one of the two tests.

### Neural activity bootstrapping

We used bootstrapping methods to compute percentiles and p-values to quantify the responsiveness of neurons to saccades. We used dF/F traces provided by the Allen Brain Observatory (de Vries et al., 2020). For every neuron, we computed the average dF/F in frames 0 to 10 (0ms to 333ms) relative to the saccade onset for each saccade, along with the average dF/F in frames −45 to −15 (−1.5s to −0.5s) relative to the saccade onset. We then subtracted the latter from the former, yielding the change in dF/F value for each neuron and saccade. We computed the mean change in dF/F for each neuron, referred to as its mean saccade response, by averaging the values across all saccades.

We used this same response computation to build a bootstrapped baseline distribution of containing *N* = 40,000 samples for each neuron, where each sample was computed using the same procedure above, except the onset times was determined using the following procedure: (i) randomly choose an experiment session among the set of sessions in which a neuron was present, where the sessions are weighted by their total number of saccades; (ii) select a frame uniformly at random across the entire chosen session. We then computed the quantile *q* (expressed as a value between 0 and 1) of the neuron’s mean saccade response within the bootstrap distribution. Specifically, if *q* was within *p* of 1, we flagged the neuron as having an enhanced response to saccades; if *q* was within *p* of 0, we flagged the neuron as having a suppressed response to saccades; and otherwise we flagged the neuron as having no response to saccades. Saccade responsive (SR) neurons were those that had either an enhanced or suppressed response to saccades, using *p* = 0.05, corrected for multiple comparisons.

### Direction selective inclusion criteria

Using an analogous procedure as in (Miura & Scanziani, 2022), we compared the average dF/F in frames 0 to 10 relative to the saccade onset in nasal and temporal directions using a Wilcoxon rank-sum test, and flagged a SR neuron as direction selective (DS) if the result of this test was significant (p < 0.05). For DS neurons, we defined the preferred direction as the direction (nasal or temporal) that maximized the neuron’s mean saccade response to saccades along that direction.

### Saccade neural responses to different visual stimuli

To see how a saccade-responsive neuron’s response varied depending on different stimuli, we first considered the neuron’s saccade response classification: if the neuron was responsive to saccades in a direction-selective manner, then we considered only saccades in the preferred direction; otherwise, we considered saccades in all directions. We excluded experimental sessions containing fewer than 8 saccades during the spontaneous stimulus. For each neuron, we computed the response to each saccade using the same procedure as in Methods, “Neural activity bootstrapping.” We used a Wilcoxon rank-sum test on related paired samples (namely, the mean response to saccades made during spontaneous and the mean response to saccades made during the other compared visual stimulus). We computed normalized responses for each session by dividing the neuron’s mean response for saccades made during each stimulus by its mean response to saccades made during the spontaneous stimulus.

### Visual response metrics

To assess the visual responses of SR neurons, we used response metrics previously computed for the dataset and described in (de Vries et al., 2020). For each neuron’s response to drifting gratings, the preferred condition was identified as the grating direction and temporal frequency at which the neuron had its largest mean response. This preferred condition determined the neuron’s preferred direction.

Direction selectivity index (DSI) for each neuron was computed using its mean response to the drifting gratings stimulus at its preferred temporal frequency:

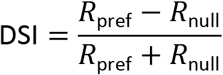

where *R*_pref_ is the neuron’s mean response to its preferred direction and *R*_null_ is its mean response to the opposite direction at the same temporal frequency.

Orientation selectivity index (OSI) was computed from the mean response to the drifting gratings stimulus at each neuron’s preferred temporal frequency as

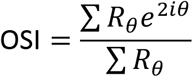

where *R_θ_* is the neuron’s mean response to direction *θ* (Ringach et al., 1997).

The lifetime sparseness was computed using the definition in (Vinje & Gallant, 2000), as

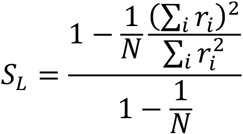

where *N* is the number of stimulus conditions and *r_i_* is the response of the neuron to stimulus condition *i* averaged across trials.

## Supplementary Figures

**Figure 3, Supplement 1:**
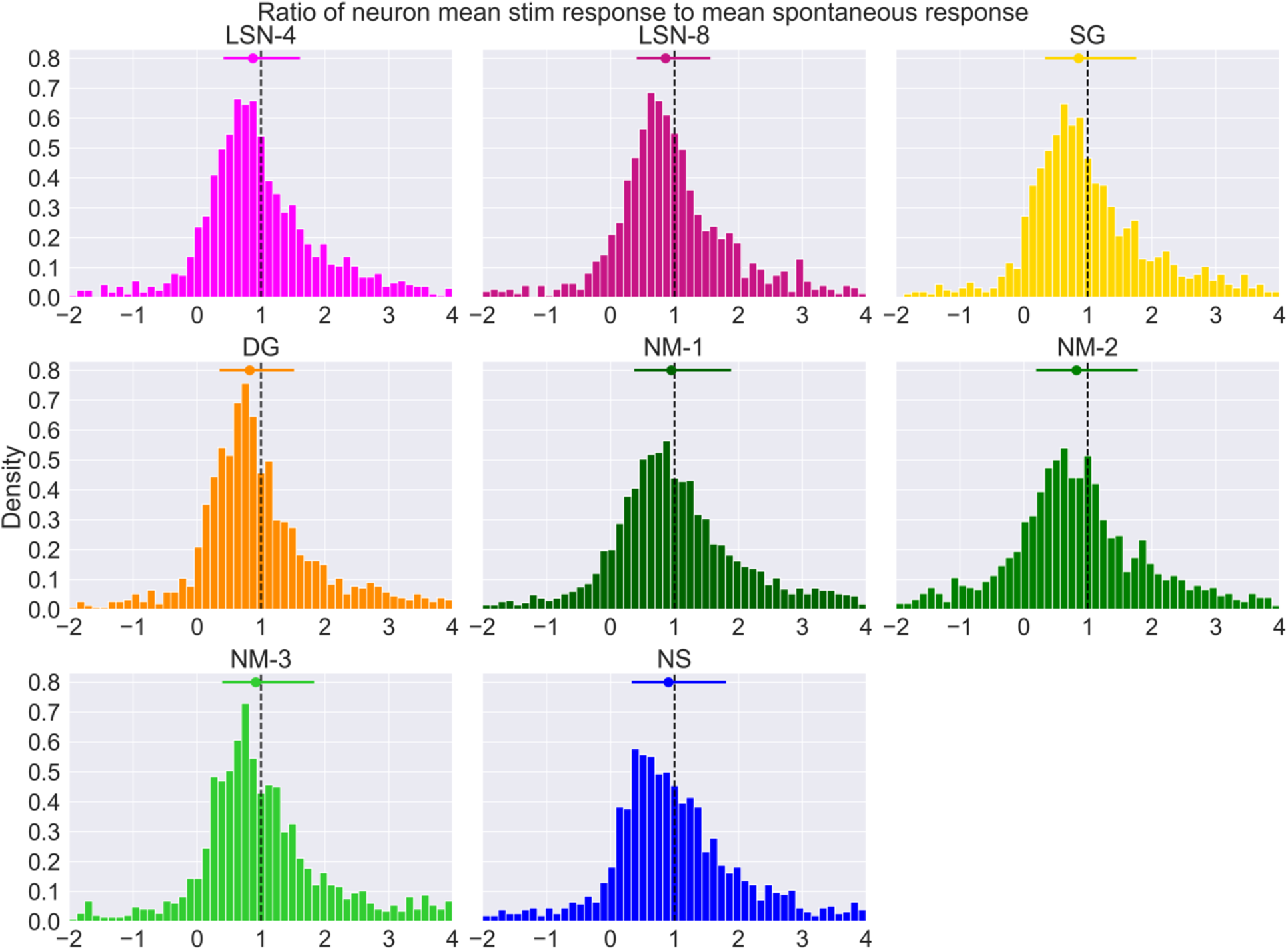
Ratio of mean saccade response during each stimulus to mean response during gray screen. Dashed vertical line at x=1 indicates equal mean response. Top bar and dot represent middle 50% and median, respectively.

**Figure 4, Supplement 1:**
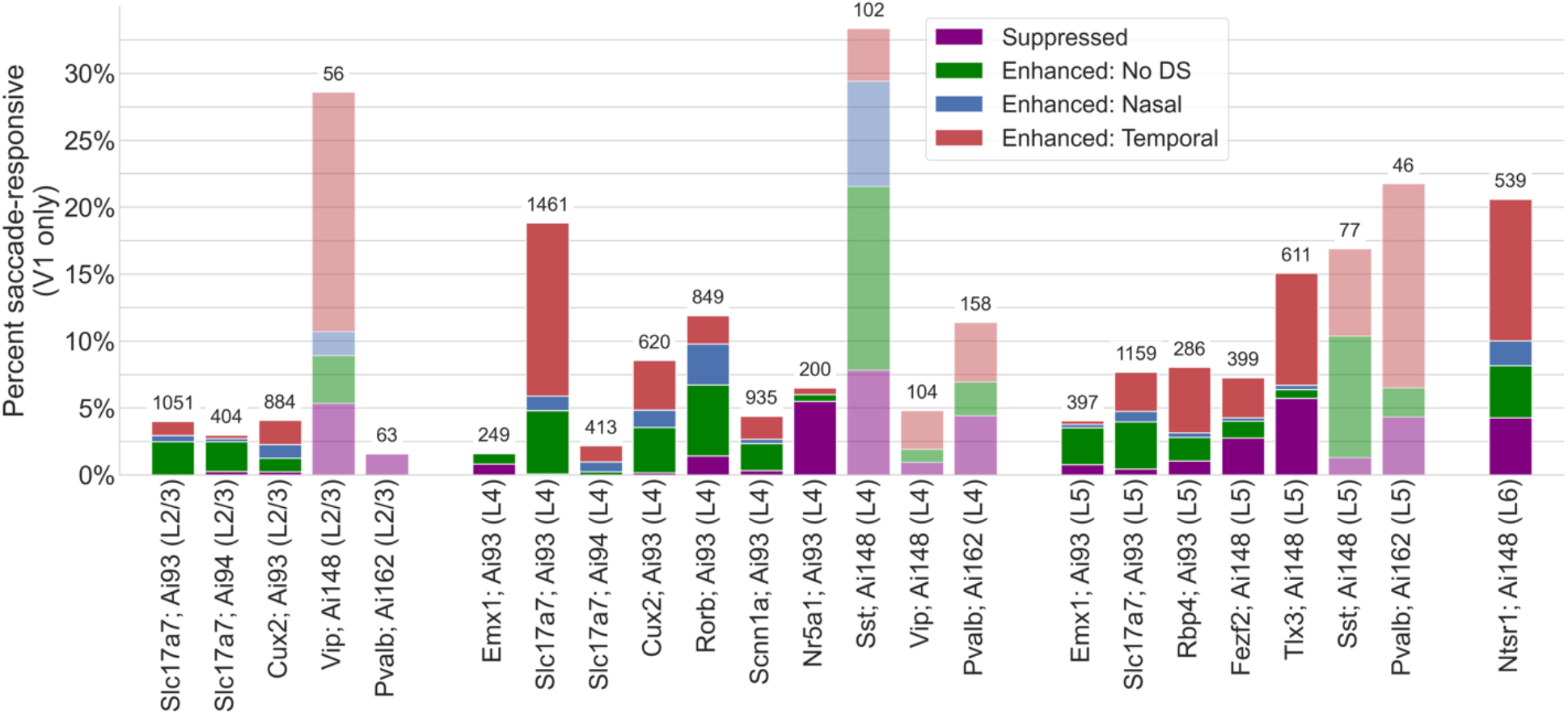
Percent of saccade responsive neurons in V1. Percent SR neurons by transgenic line and cortical layer, across only V1. Low opacity bars indicate inhibitory neurons, and numbers above bars indicate the total number of neurons imaged.

